# Heparan sulfate structure is influenced by the ER-Golgi dynamics of its modifying enzymes

**DOI:** 10.1101/2020.01.23.916940

**Authors:** Maria Cecília Zorél Meneghetti, Paula Deboni, Carlos Modesto Vera Palomino, Luiz Patekoski Braga, Renan Pelluzzi Cavalheiro, Gustavo Monteiro Viana, Edwin A. Yates, Helena B. Nader, Marcelo A. Lima

**Author notes:** Corresponding author (M A. Lima).

## Abstract

The cell surface and extracellular matrix polysaccharide, heparan sulfate (HS) conveys chemical information to control or influence crucial biological processes. Attempts to describe its structure-function relationships with HS binding proteins in a classical ‘lock and key’ type manner, however, have been unsuccessful. HS chains are synthesized in a non-template driven process in the ER and Golgi apparatus, involving a large number of enzymes capable of fine-tuning structures. Changes in the localization of HS-modifying enzymes throughout the Golgi, rather than protein expression levels, were found to correlate with changes in the structure of HS. Following brefeldin A treatment, the HS-modifying enzymes localized preferentially in COPII vesicles and at the trans-Golgi. Further, shortly after treatment with heparin, the HS-modifying enzyme moved from cis to trans-Golgi, which coincided with increased HS trisulfated disaccharide content. Finally, it was shown that COPI subunits and Sec24 gene expression changed. Collectively, these findings highlight that the ER-Golgi dynamics of HS-modifying enzymes via vesicular trafficking processes are critical prerequisite for the complete delineation of HS biosynthesis.

## Introduction

Protein glycosylation, the post-translational modification of proteins in which carbohydrate moieties are conveniently attached, is seen as the new frontier in the field of glycomics (Martin et al., 2009). Glycosylation is the most abundant post-translational modification and plays a vital role in protein function (Haltiwanger and Lowe, 2004). Heparan sulfate (HS) is a sulfated glycosaminoglycan (GAG) found on the cell membrane and in the extracellular matrix throughout the animal kingdom (Cássaro and Dietrich, 1977; Medeiros et al., 2000). Alongside heparin (Hep), HS is a member of the GAG family which are present in tissues as proteoglycans, where the polysaccharide chains are covalently bound to a protein backbone. Their chains are mainly composed of repeating disaccharide units of 1,4 linked uronate, either β–D-glucuronate or α-L-iduronate, and α-D-glucosamine, where N-acetyl-D-glucosamine residues become de-N-acetylated and N-sulfated, then, β–D-glucuronates undergo epimerization at C5 to α-L-iduronates (IdoA). Furthermore, sulfate groups may be added at C2 of uronate residues, C6 of the glucosamine residues, and less commonly, at C3 of glucosamine residues (Dietrich et al., 1983; Meneghetti et al., 2015). Such fine-tuning is the result of a series of enzymatic reactions that do not modify the HS chains completely, giving rise to complex substitution patterns.

A central hypothesis in the field is that the HS chain substitution pattern encodes its capability to influence many key biological processes (Cavalheiro et al., 2017; Moreira et al., 2004; Nader et al., 1999; Sarrazin et al., 2011), through interactions with hundreds of proteins (Nunes et al., 2019). Yet, initial attempts to understand structure-function relationships in the classic ‘lock and key’ mechanism were almost entirely unsuccessful even in the well-studied examples of FGF:FGFR signaling and AT activity (Guerrini et al., 2014; Kan et al., 1996; Lima et al., 2013; Mohammadi et al., 2005; Rudd et al., 2010; Xu et al., 2012; Xu et al., 2013). It is now appreciated that there exists complex and regulated biosynthetic machinery capable of producing finely-tuned HS structures and that the heterogeneity characteristic of this system will affect networks of proteins, and eventually, become evident in biological terms.

Template driven biosynthesis is employed for nucleic acids and proteins, but the biosynthesis of HS exhibits no analogous system. The order in which the enzymatic reactions are carried out by GAG modifying enzymes positioned throughout the ER and Golgi apparatus is a source of continuing debate despite several attempts to determine biosynthetic routes. A feasible order for the enzyme activity has been proposed and a biosynthetic outline able to explain the relative abundance of both common and uncommon structures and that is consistent with experimental observations has also been advanced (Meneghetti et al., 2017; Rudd and Yates, 2012). Meanwhile, the cell-defined localization of HS modifying enzymes has been questioned, since they have been shown to be located at the cell surface (Delos et al., 2018). It has also been shown that the localization of EXT1/EXT2 in distinct Golgi cisternae modulates the synthesis of HS (Chang et al., 2013), suggesting that vesicular trafficking could play an important role in the regulation of HS biosynthesis. Hence, the interrogation of cargo sorting, vesicle assembly and trafficking that takes place to deliver GAG biosynthetic enzymes throughout the ER and Golgi, is necessary for the complete description of HS biosynthesis and, hence, the success of subsequent structure and function studies.

Vesicular trafficking is mediated by protein-coated vesicles called coatomer protein complex I (COPI) and II (COPII). The COPI coat comprises a complex of seven subunits α-, β-, β’-, δ-, ε-, γ- and ζ-COP, which are biochemically grouped into cage-like (B-) and an adaptor-like (F-) subcomplexes (Arakel and Schwappach, 2018). COPI vesicles mediate retrograde transport from the Golgi and ER-Golgi intermediate compartment (ERCIG) to the ER and intra-Golgi transport in anterograde and retrograde directions (Arakel and Schwappach, 2018). The COPII coat comprises a stable complex formed by four subunits Sec23, Sec24, Sec13 and Sec31 and is implicated in anterograde transport from ER to Golgi (Jensen and Schekman, 2011). Consequently, membrane traffic between the ER and the Golgi is highly dynamic and responsible for cargo sorting and delivery among the different compartments.

Studies in both yeast and mammalian cells have identified active recycling of Golgi-resident glycosyltransferases through the ER-Golgi pathway mediated by coated vesicles (Gill et al., 2010; Liu et al., 2018; Storrie et al., 1998; Todorow et al., 2000). The different localization of these enzymes in the secretory pathway allows newly synthesized glycoconjugates to encounter glycosyltransferases in a non-uniform distribution to perform glycosylation (Emr et al., 2009; Puthenveedu and Linstedt, 2005). Mechanisms involved in glycosyltransferase retention in different compartments and within distinct Golgi cisternae may ensure the production of a wide structural variety of compounds, a key characteristic of these molecules, and may reflect the complexity of glycoconjugate synthesis.

It has also become clear that the search for the precise control of HS biosynthesis through the modulation of individual enzymes is ineffective, while the Golgi dynamics remain poorly understood and the different cellular contexts encountered have been largely ignored. Artificial Golgi systems have been built as test beds to better understand how the natural Golgi controls the biosynthesis of GAGs and ultimately, for the design of bioengineered heparin (Martin et al., 2009) but, again, the natural Golgi dynamics and cellular context has not been taken into account. Thus, it seems probable that further regulatory mechanisms are at work; ones that are, perhaps, not apparent at the level of individual biosynthetic enzymes. It was with this in mind that the authors have embarked on the present investigation.

In the present study, the influence of vesicular trafficking mediated by COPI and COPII in the distribution, along the early secretory pathway, of HS-modifying enzymes in the regulation of HS biosynthesis was evaluated. Furthermore, the effect of pharmacological agents that inhibit vesicular trafficking and overstimulate HS synthesis, which leads to changes in HS-modifying enzyme dynamics and result in altered HS structure, were explored. This study represents a first step towards gaining an understanding of how the natural Golgi controls the biosynthesis of HS and to determining whether cells can be bioengineered to produce tailored GAG structures.

## Results

### Subcellular localization of fluorescently tagged HS3ST5

Initially, the distribution profile of HS-modifying enzymes in coated vesicles was analyzed in endothelial cells (EC) by confocal microscopy (Fig. S1). Regardless of the position in the hierarchical sequence of the biosynthetic process, the enzymes involved in HS biosynthesis (NDST1, C5-epimerase, HS3ST1 and HS3ST3A) showed similar localization in COPI and COPII vesicles. 3-O-sulfotransferase, the last enzyme to modify the HS chains according to the classic HS biosynthetic pathway, was chosen as a tracker and had its subcellular localization investigated using overexpression systems in EC cells. The overexpression of this HS-modifying enzyme was confirmed by RT-PCR using cloning primers (HS3ST5-BglII and HS3ST5-BamHI), in concert with confocal microscopy and flow cytometry. Only transfected cells showed a band at the expected size for tagged-HS3ST5 (Fig. S2 A). In addition, EC-HS3ST5 cells also showed increased total and tagged HS3ST5 protein expression levels, which were analyzed by anti-HS3ST5 antibody (Fig. S2 B) and direct fluorescence of GFP assays (Fig. S2 C), respectively.

The localization of the overexpressed protein in the Golgi apparatus and coated vesicles was confirmed by immunostaining of the cells using specific antibodies labelled with Alexa Fluor^®^ 633. As shown in Fig. 1 A, HS3ST5 colocalized both with GM130, cis-Golgi protein marker, and Golgin97, a trans-Golgi protein marker, confirming HS3ST5 localization in both cisternae of the Golgi apparatus and highlighting that the enzyme is trafficked continually amongst the different Golgi cisternae. HS3ST5 exhibited a similar distribution in COPI, exemplified by α-COP and β-COP subunits, and COPII, represented by staining of the Sec23 subunit, further confirming that HS3ST5 is cycled constantly in the ER-Golgi pathway (Fig. 1 B). Further analysis has been conducted using super resolution microscopy. The results clearly showed the colocalization and putative interface of HS3ST5 with different coated vesicle subunits (Fig. 1 C).

**Figure 1.**
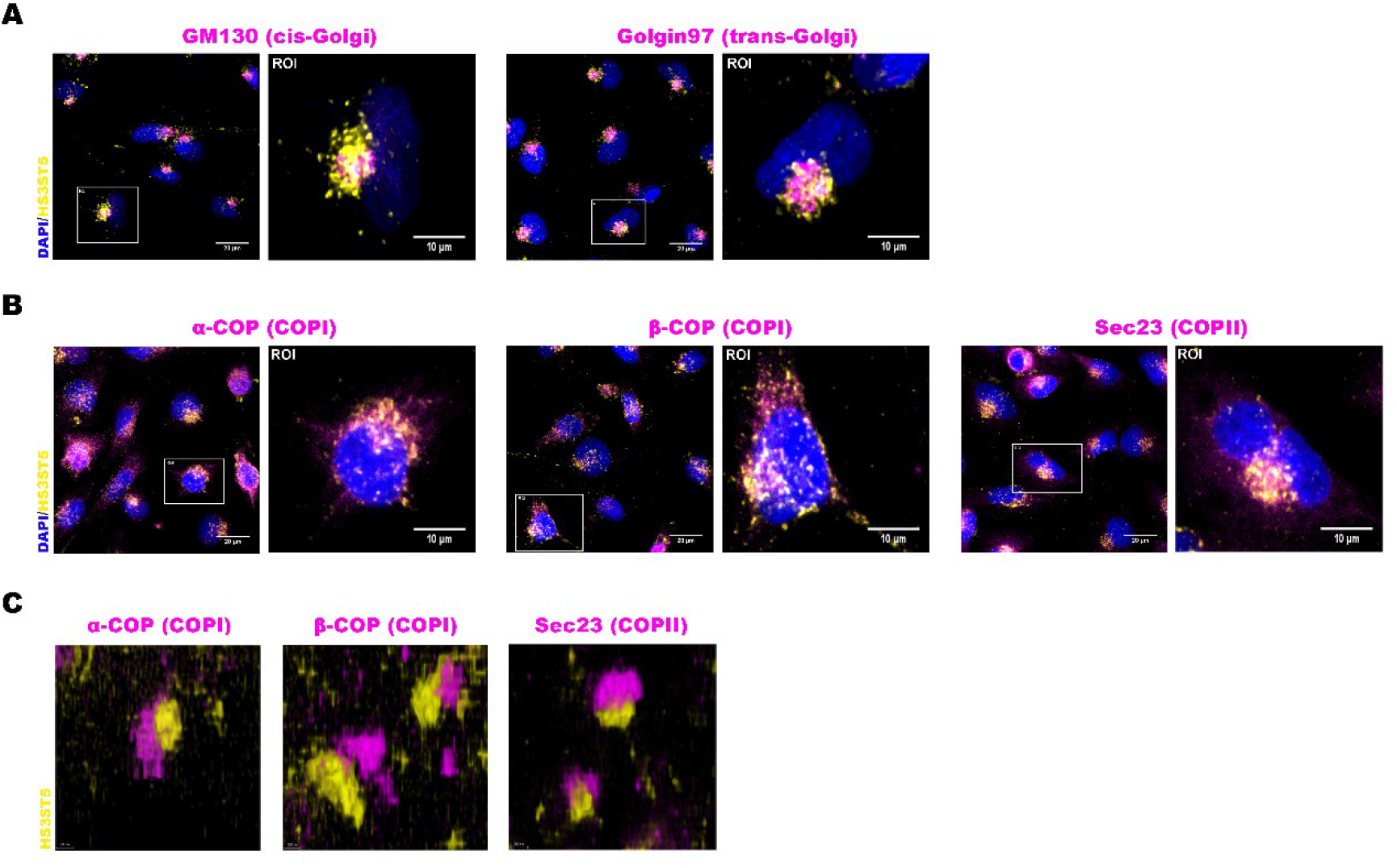
Subcellular localization of fluorescent HS3ST5. **(A)** Transfected cells were labeled with anti-GFP antibody (tagged HS3ST5) and specific antibodies to cis-Golgi (GM130) or trans-Golgi (Golgin97). The staining was revealed with secondary antibodies conjugated with Alexa Fluor® 488 (yellow) and Alexa Fluor® 633 (magenta), respectively. Fluorescent images were analyzed on TCS SP8 CARS confocal microscopy. **(B)** EC-HS3ST5 were doubled-labelled with antibodies to GFP and to coated vesicles and revealed with secondary antibodies conjugated with Alexa Fluor® 488 (yellow) and Alexa Fluor® 633 (magenta), respectively. COPI vesicles were visualized by α-COP and β-COP staining and COPII vesicles were visualized by Sec23 staining. **(C)** Super resolution microscopy images of HS3ST5 and COPI and COPII vesicles. The transfected cells were labeled with anti-GFP and specific antibodies to coated vesicles, revealed with secondary antibodies conjugated Alexa Fluor® 647 (yellow) and Alexa Fluor® 488 (magenta) respectively and analyzed on SR-GSD microscopy. Representative microscopy images of expression of recombinant HS3ST5 and coated vesicles or Golgi markers are shown in yellow and magenta staining, respectively. Nuclei were stained with DAPI (blue). Scale bars in images: 20 μm; Scale bars in ROI images: 10 μm. ROI: region of interest.

### Effects of brefeldin A on the localization of HS-modifying enzymes in vesicular trafficking

In order to evaluate the influence of vesicular trafficking on the transport of HS-modifying enzymes along the secretory pathway, EC-HS3ST5 cells were treated with brefeldin A (BFA), a pharmacological inhibitor of ADP-ribosylation factors, which are responsible for recruitment of COPI subunits (Peyroche et al., 1999). In the presence of BFA, HS3ST5 displayed higher levels of colocalization in COPII vesicles, indicating that the enzyme was maintained during anterograde transport (Fig. 2 A). It is known that BFA causes Golgi cisternae disassembly and the redistribution of proteins from the cis and medial-Golgi into the ER (Kolset et al., 2002; Lippincott-Schwartz et al., 1989). As expected, the BFA treatment induced disassembly of the Golgi indicated by GM130 and Golgin97 scattered staining (Fig. 2 B). The effect of BFA was also followed by changes in HS3ST5 distribution along the Golgi cisternae from cis- to trans-Golgi (Fig. 2 B).

**Figure 2.**
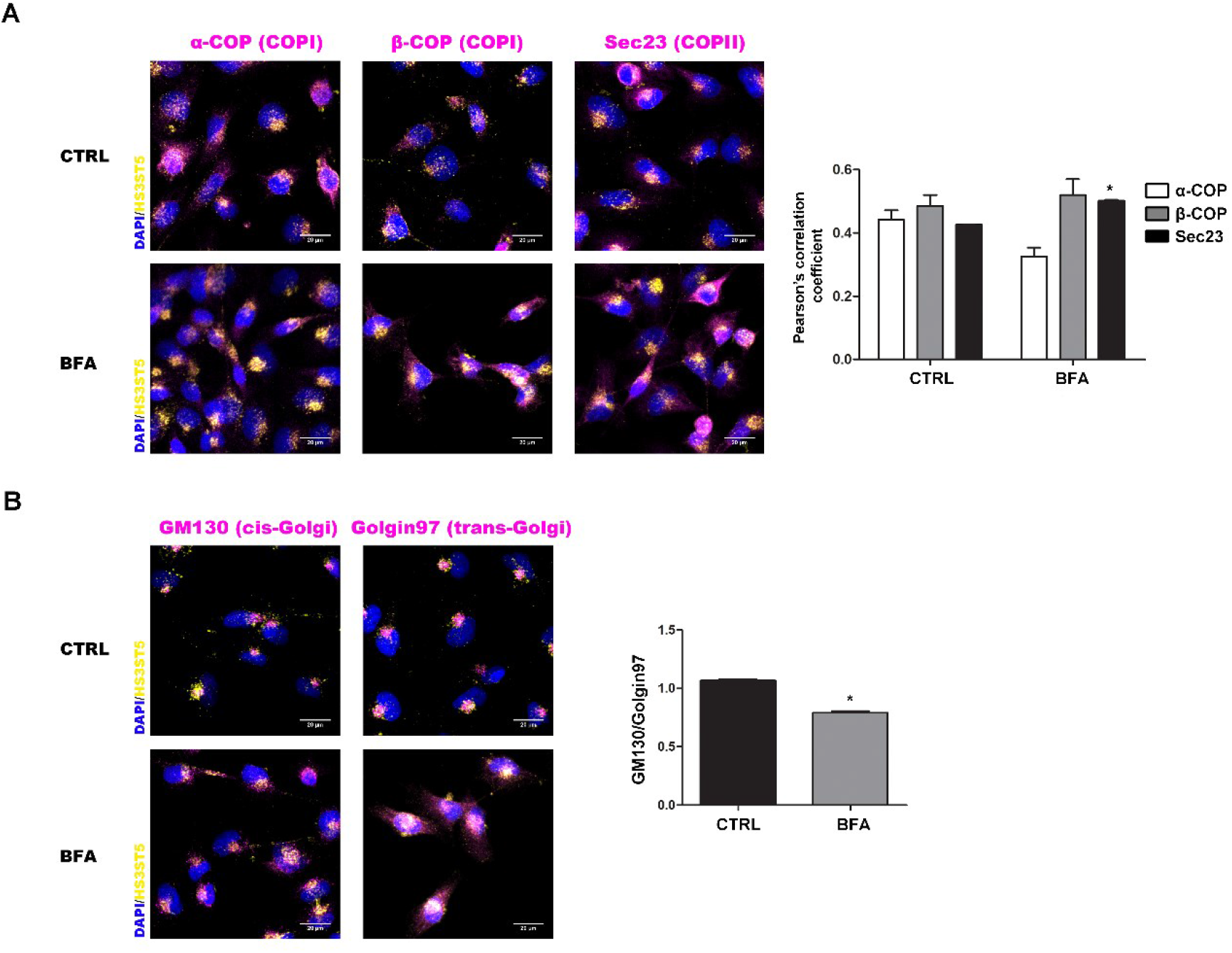
Distribution profile of HS3ST5 in secretory pathway in the presence of brefeldin A. **(A)** After treatment with BFA (3 µg/mL) for 2h, EC-HS3ST5 cells were labeled with anti-GFP (tagged HS3ST5) and specific primary antibodies to coated vesicles and then revealed with secondary antibodies conjugated with Alexa Fluor® 488 (yellow) and Alexa Fluor® 633 (magenta), respectively. Fluorescent images were analyzed on confocal microscopy. Pearson’s correlation coefficient represents rate of colocalization of recombinant HS3ST5 in coated vesicles and was obtained using the Leica LAS X Life Science software (right panel). COPI vesicles were visualized by α-COP and β-COP staining and COPII vesicles were visualized by Sec23 staining. **(B)** EC-HS3ST5 previously treatment with BFA were double-labelled with anti-GFP antibody and specific antibodies to GM130 (cis-Golgi) or Golgin97 (trans-Golgi) and then revealed with secondary antibodies conjugated with Alexa Fluor® 488 (yellow) and Alexa Fluor® 633 (magenta), respectively. Right panel corresponds to the ratio of Pearson’s correlation coefficients obtained of recombinant HS3ST5 in cis-Golgi (GM130) and in trans-Golgi (Golgin97). Nuclei were stained with DAPI (blue). Scale bars in images: 20 μm. *, Differences statistically significant, *P < 0.05*, relative to control.

The profile of other HS-modifying enzymes (NDST1, C5-epimerase, HS2ST, HS3ST1 and HS3ST3A) in the presence of BFA, relative to HS3ST5, was also analyzed by immunostaining and confocal microscopy. All enzymes presented colocalization with HS3ST5 (Fig. 3), showing that the HS-modifying enzymes are localized in the same Golgi cisternae.

**Figure 3.**
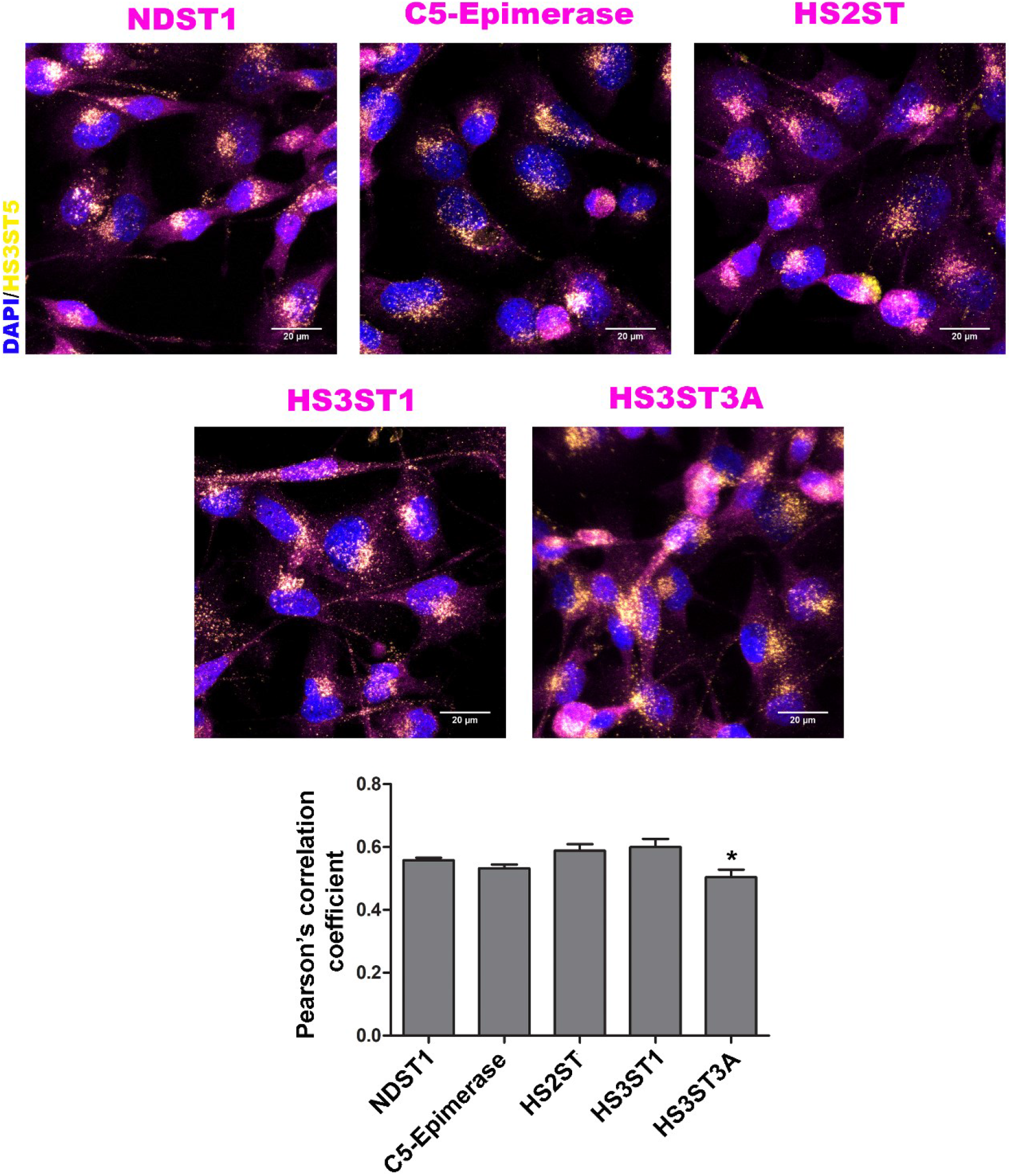
Distribution profile of HS-modifying enzymes following BFA treatment. After treatment with BFA (3 µg/mL) for 2h, EC-HS3ST5 cells were double-labeled for HS3ST5-GFP and HS-modifying enzymes (NDST1, C5-Epimerase, HS2ST, HS3ST1 and HS3ST3A). Secondary antibodies conjugated with Alexa Fluor® 488 (yellow) and Alexa Fluor® 633 (magenta), respectively were used. Fluorescent images were acquired and analyzed by confocal microscopy. Tagged HS3ST5 and other HS-modifying enzymes are shown in yellow and magenta staining, respectively. The nuclei (blue) were stained with DAPI. Pearson’s correlation coefficient represents the rate of colocalization of the tagged HS3ST5 with each HS-modifying enzyme. Scale bars in images: 20 μm. *, Statistically significant, *P < 0.05*, relative to HS3ST1.

### Vesicular trafficking and Golgi apparatus localization of HS3ST5 changes with heparin treatment

Endothelial cells in the presence of heparin upregulate the synthesis of HS with increasing amounts of trisulfated disaccharide (Nader et al., 1989). In order to assess whether this was the result of changes in the localization of HS-modifying enzymes along the Golgi cisternae, the cells were exposed to heparin and the distribution profile of the HS3ST5 relative to coated vesicles and Golgi apparatus was analyzed by confocal microscopy after immunofluorescence staining. There were no changes in HS3ST5 distribution in either COPI and COPII vesicles (Fig. 4). However, changes in HS3ST5 distribution within the Golgi were observed (Fig. 5). While the HS3ST5 was preferentially present in the cis-Golgi in cells with no treatment, or during the first hour of heparin exposure, HS3ST5 changed its distribution from cis to trans-Golgi in subsequent hours (2-3 h).

**Figure 4.**
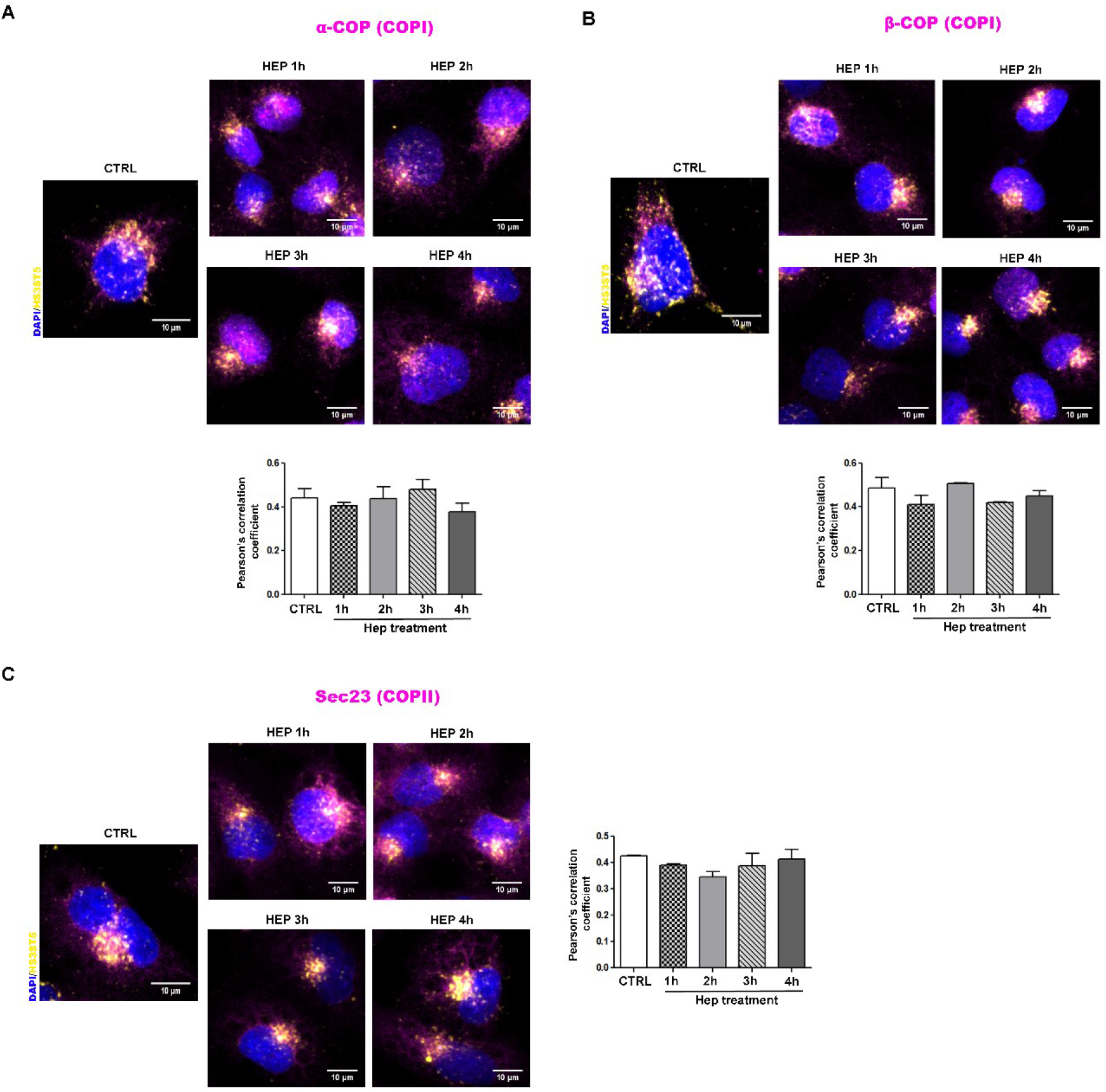
Distribution profile of HS3ST5 in coated vesicles in the presence of heparin. After treatment with heparin (20 µg/mL) from 1 to 4 h, EC-HS3ST5 cells were double-labeled with antibodies to GFP (tagged HS3ST5) and α-COP **(A)**, β-COP **(B)** or Sec23 **(C)**. The cells were revealed with secondary antibodies conjugated with Alexa Fluor® 488 or Alexa Fluor® 633. Recombinant HS3ST5 and coated vesicles are shown in yellow and magenta staining, respectively. The nuclei (blue) were stained with DAPI. Pearson’s correlation coefficient represents rate of colocalization of recombinant HS3ST5 in coated vesicles (down and right panels). Scale bars in images: 10 μm.

**Figure 5.**
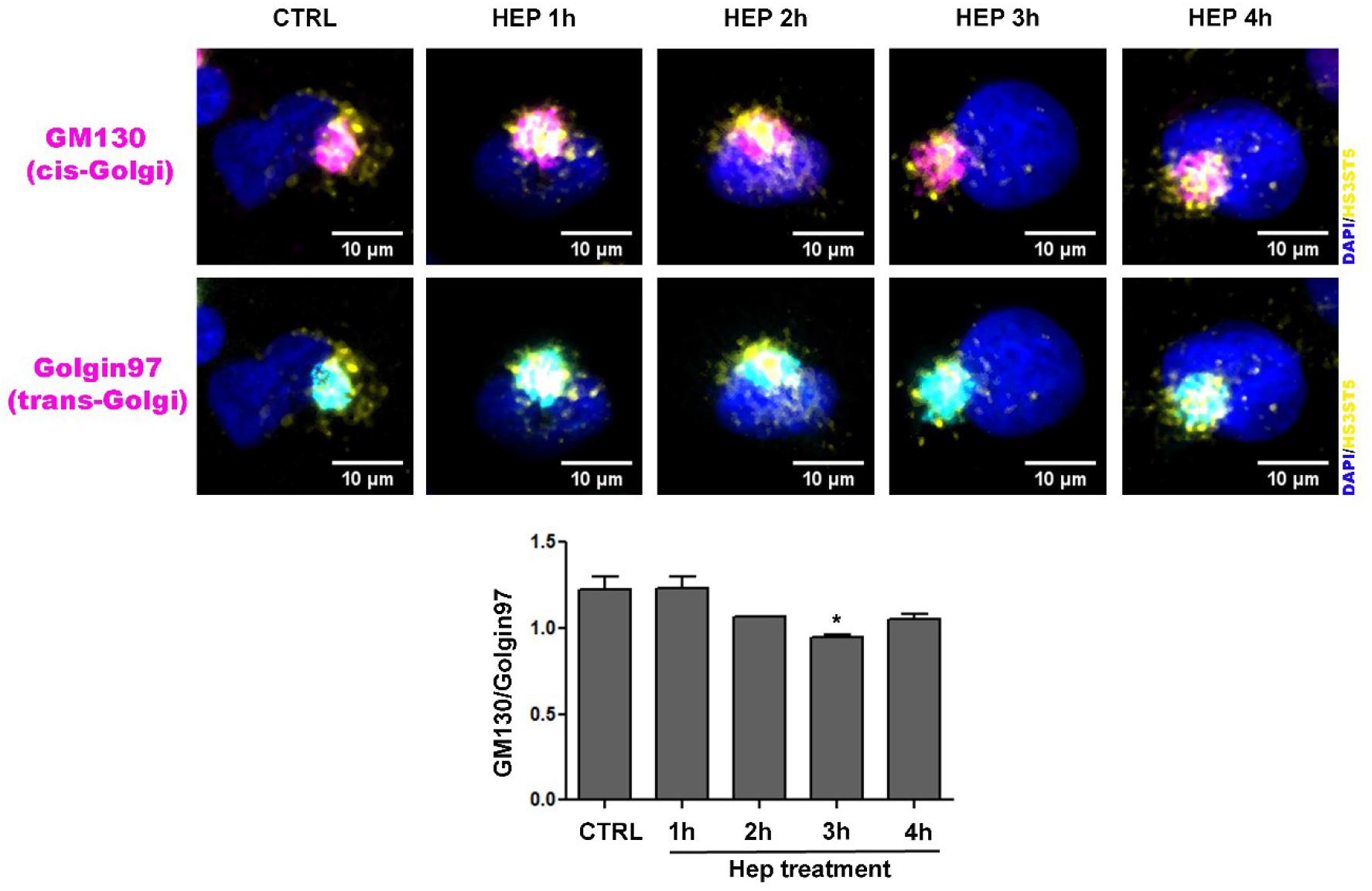
Distribution profile of HS3ST5 in Golgi apparatus following heparin treatment. After treatment with heparin (20 µg/mL) for 1 to 4 h, EC-HS3ST5 cells were triple-staining for GFP (tagged HS3ST5), GM130 (cis-Golgi) and Golgin97 (trans-Golgi). Secondary antibodies conjugated with Alexa Fluor® 488, Alexa Fluor® 594 and Alexa Fluor® 633, respectively were used. Tagged HS3ST5 is show in yellow, whereas GM130 and Golgin97 are shown in magenta and cyan, respectively. The nuclei (blue) were stained with DAPI. The ratio GM130/Golgin97 corresponds to the Pearson’s correlation coefficients obtained for the tagged HS3ST5 in the cis-Golgi and in the trans-Golgi, respectively (bottom panel). Scale bars in images: 10 μm. *, Statistically significant, *P < 0.05*, relative to control.

### Vesicular trafficking of HS3ST5 influences the structure of HS

In order to confirm the functional relationships between the structure of HS and vesicular trafficking under heparin stimulation, we performed metabolic ^35^S-sulfate labeling of HS followed by analysis of the resulting HS structure by enzymatic digestion. The disaccharide composition revealed an increase in the trisulfated disaccharide proportion for the HS present in the cell extract and secreted to the media (Fig. 6 and Table S1). As expected, changes were first detected in the cell extract (Fig. 6 A) followed by changes in the HS secreted to the media (Fig. 6 B). Importantly, these results correlate with the changes in HS3ST5 trafficking from cis- to trans-Golgi and recycling to cis again (Fig. 5). Together, they show that vesicular trafficking regulates the sulfation pattern found in HS.

**Figure 6.**
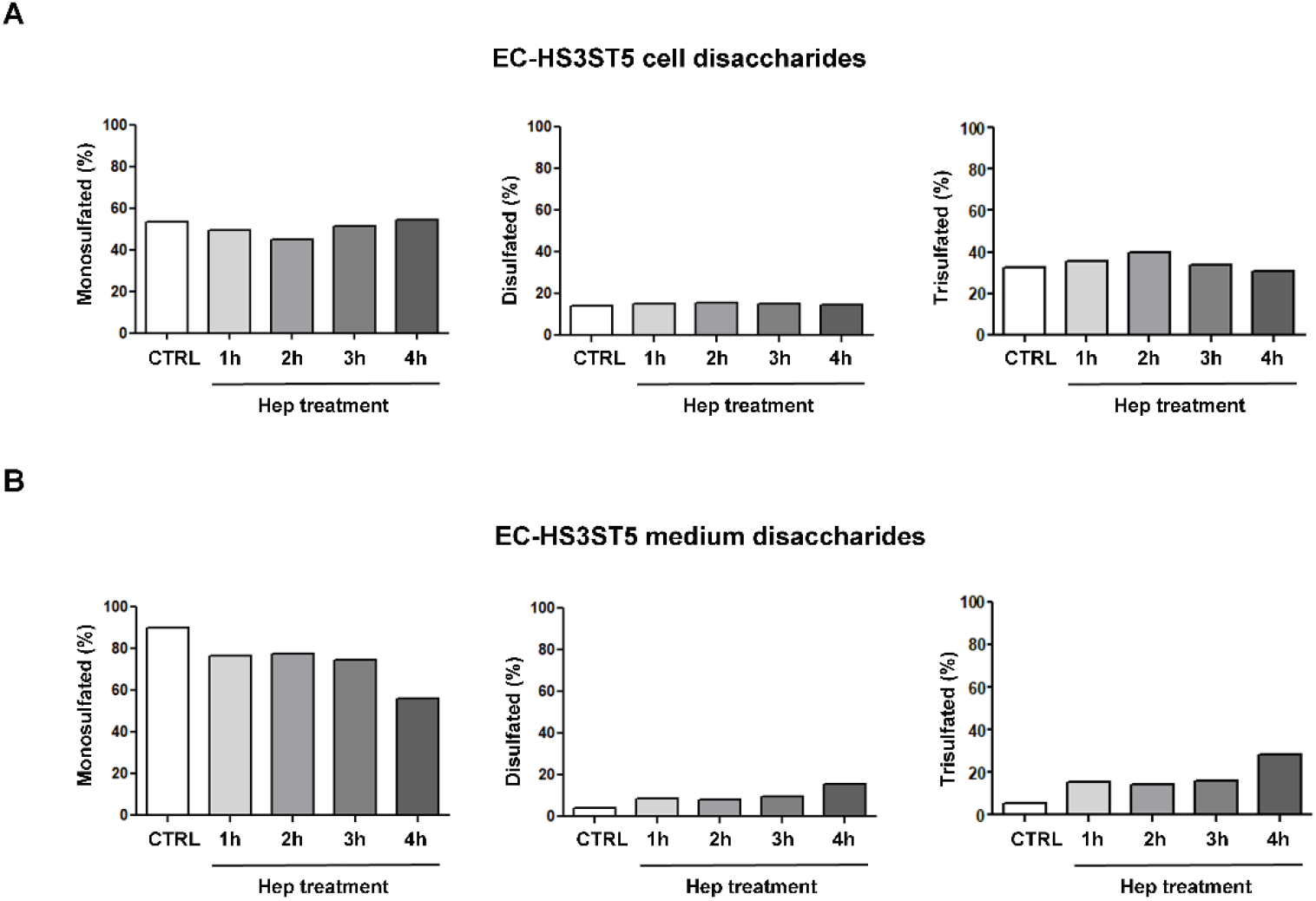
Disaccharide composition of HS extracted from EC-HS3ST5 cells treated with heparin. After treatment with heparin (20 µg/mL) from 1 to 4 h, GAGs labeled with [^35^S]H2SO4 were purified from the EC-HS3ST5 cells **(A)** and culture medium **(B)** as described in methods. The samples were digested using heparitinases I and II from *Flavobacterium heparinum* and Δ-disaccharides of HS were analyzed in ion exchange liquid chromatography. Individual fractions (0.3 mL) were collected and counted on a Micro-Beta counter. The Δ-disaccharides of HS were shared in monosulfated, disulfated and trisulfated groups. The values are representative of a pool of three independent experiments.

### HS-modifying enzymes and PAPS synthase levels are not up-regulated following heparin treatment

The increase in HS sulfation could also be the result of sulfotransferase along with PAPS synthase upregulation. To further confirm our hypothesis that the vesicle trafficking instead is responsible for the detected structural changes, protein and gene expression experiments were conducted. Flow cytometry analysis for specific HS-modifying enzymes (NDST1, C5-Epimerase, HS2ST and HS3ST5) indicated that the protein levels remained unchanged throughout the heparin treatment (Fig. 7 A). Gene expression analysis also showed significant decrease in both PAPS synthase isoforms during heparin treatment (Fig. 7 B). Together, these results further support the hypothesis that vesicular trafficking, rather than protein/gene expression, regulates HS biosynthesis.

**Figure 7.**
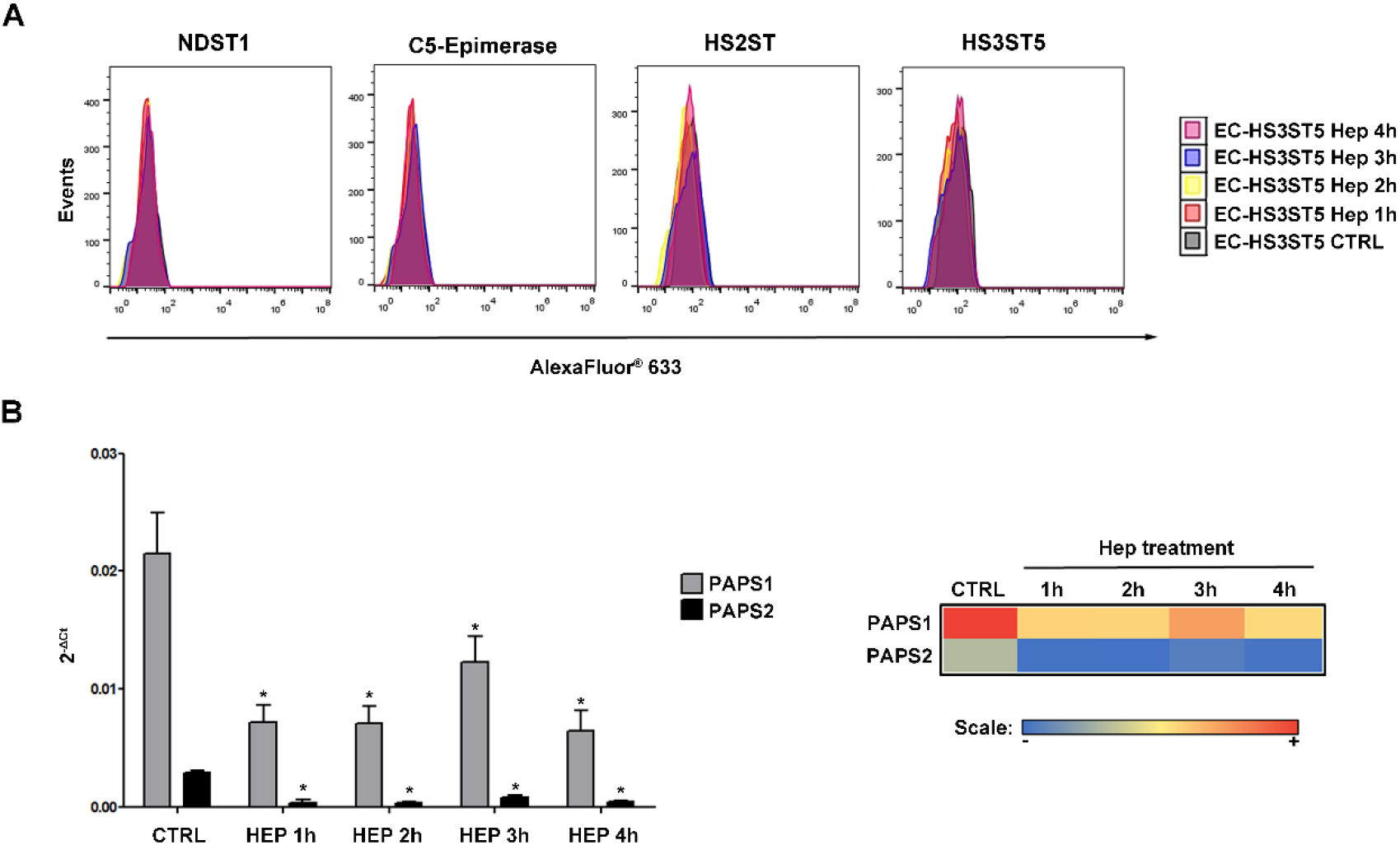
Protein and gene expression of components of HS biosynthesis in presence of heparin. **(A)** Protein expression of HS-modifying enzymes (NDST1, C5-epimerase, HS2ST and HS3ST5) in transfected cells previously treated with heparin was evaluated by flow cytometry using antibodies specific for each enzyme. Following incubation with primary antibodies, cells were incubated with secondary antibody conjugated with Alexa Fluor® 633 and analyzed on FACSCalibur flow cytometer. **(B)** PAPS synthases mRNA level in EC-HS3ST5 cells treated with heparin was analyzed by real-time. The results were expressed as mean ± standard deviation. (Right panel) Heat map was generated of mean values obtained in the gene expression assays. High and low expression are shown in red and blue colors. *, Differences statistically significant, *P < 0.05*, relative to control.

### Changes in coated vesicle component expression after heparin stimulus

Finally, gene expression analysis of COPI subunits as well as Sec24 subunit isoforms of COPII during heparin stimulation were performed in order to evaluate the relationship between trafficking of HS-modifying enzymes and the expression of coated vesicle subunits responsible for cargo binding and sorting. It is known that while all seven COPI subunits are engaged in cargo recognition (Arakel and Schwappach, 2018; Watson et al., 2004; Yu et al., 2009), multiple isoforms of Sec24 are the major cargo binding subunit within the COPII vesicle (Mancias and Goldberg, 2008; Miller et al., 2002). Fig. 8 A shows that the gene expression for most COPI subunits comprising both B- and F-subcomplexes, changed after stimulation with heparin. Compared to the controls, β’-COP, β-COP and δ-COP subunits showed reduced gene expression in the early stages of treatment, while gene expression of γ1-COP, γ2-COP and ζ1-COP only changed later. Whereas the γ-COP1 subunit showed a reduction in its mRNA level in 4 h, γ2-COP and ζ1-COP presented significant increases in their gene expression during this time period; ζ-COP1 being the principal COPI subunit experiencing the highest modification in gene expression. As for Sec24, gene expression of only isoforms A and B changed, and reduction in mRNA levels alone during the early phase of heparin stimulus was observed (Fig. 8 B).

**Figure 8.**
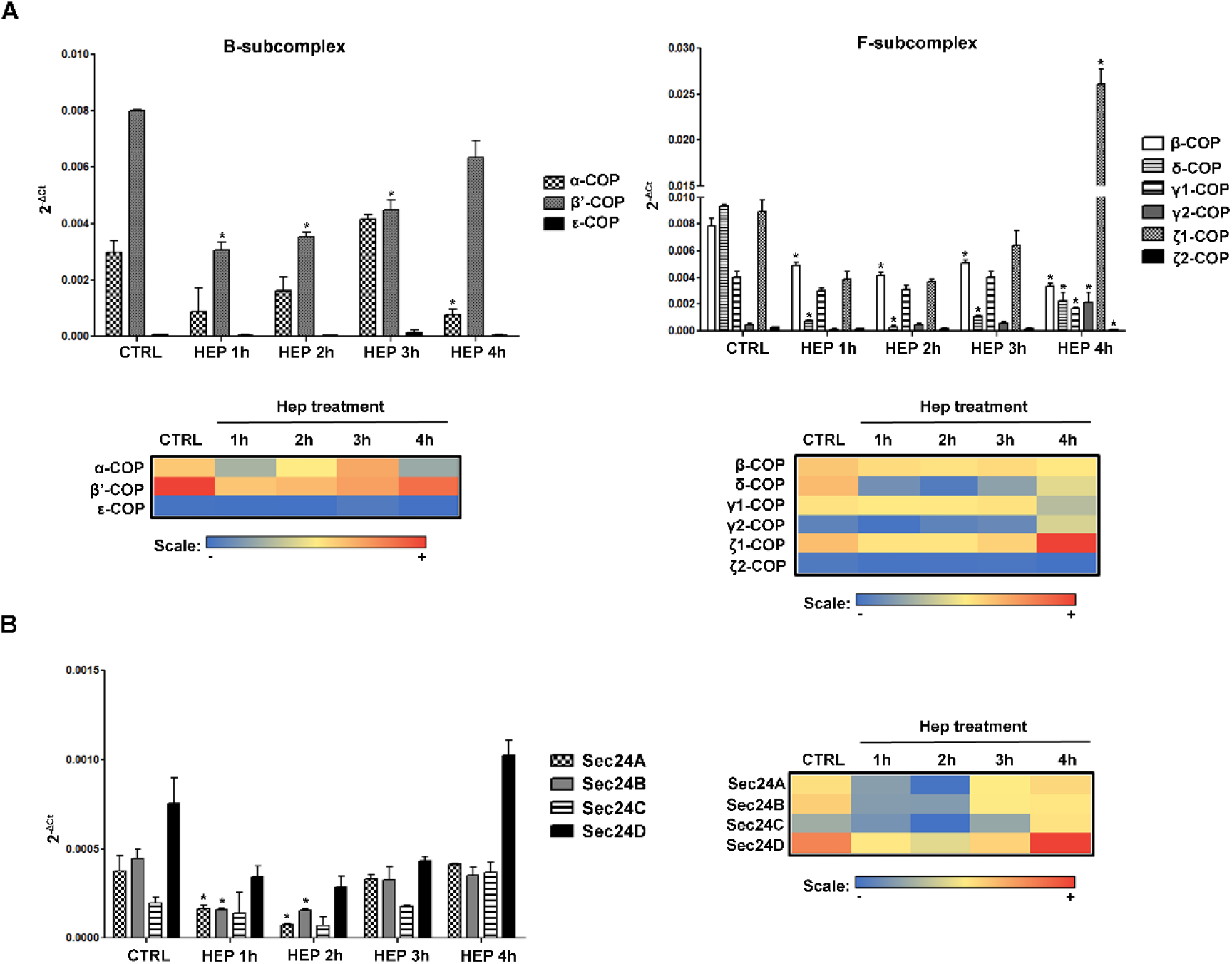
Gene expression of coated vesicles subunits in the presence of heparin. COPI subunits subdivided in B- and F-subcomplex **(A)** and Sec24 subunit **(B)** mRNA level in EC-HS3ST5 cells treated with heparin was analyzed by real-time. The results were expressed as mean ± standard deviation. Heat maps were generated of mean values obtained in the gene expression assays. High and low expression are shown in red and blue colors. *, Differences statistically significant, *P < 0.05*, relative to control.

In summary, the results show that upon heparin treatment, cargo sorting associated proteins have their gene expression altered first, followed by changes in genes that code for coat proteins linked to vesicle trafficking within the Golgi cisternae. Collectively, these results are in agreement with the spatial and temporal changes observed in the Golgi distribution of HS-modifying enzymes that preceded the biosynthesis of HS with increasing amounts of trisulfated disaccharide.

## Discussion

The structural diversity of HS could conceivably arise from many cellular events that regulate HS biosynthesis and, consequently, influence HS substitution pattern. The structural variability could, therefore, have been due to UDP-sugar and PAPS availability in Golgi cisternae (Dick et al., 2012), the interaction among HS-modifying enzymes themselves and among other proteins (Fang et al., 2016; Pinhal et al., 2001; Presto et al., 2008; Senay et al., 2000), as well as their availability and distribution throughout the ER and Golgi. It has been shown, however, that the vesicular trafficking influences both spatial and temporal localization of many glycosyltransferases along the ER-Golgi pathway, regulating the sequential order in which these enzymes act upon glycoconjugate synthesis (Tu and Banfield, 2010).

In the current work, we investigated the influence of COPI and COPII coated vesicles on the trafficking of HS-modifying enzymes in early secretory pathways using endothelial cells previously transfected with tagged HS3ST5. We observed that the tagged enzyme displayed similar distribution in coated vesicles, as well as in both cis- and trans-Golgi. However, the HS3ST5 distribution profile changed upon pharmacological treatment.

In the presence of BFA, HS3ST5 localized preferentially in COPII vesicles and at the trans-Golgi. Furthermore, regardless of the position in the hierarchical sequence of the biosynthetic process, other enzymes involved in HS biosynthesis (NDST1, C5-Epimerase, HS2ST, HS3ST1 and HS3ST3A) displayed similar localization, suggesting that these enzymes are also present in the trans-Golgi. Previous studies have shown that enzymes involved in the glycosylation of proteoglycans display distinct subcellular localization in rat ovarian granulosa cells in the presence of BFA and, whereas the CS/DS-modifying enzymes are exclusively distributed in the trans-Golgi, the HS-modifying enzymes are mainly located in the cis-Golgi (Uhlin-Hansen and Yanagishita, 1993). Nonetheless, other reports have also demonstrated that NDST and HS6ST are localized in the trans-Golgi of endothelial and renal epithelial cells (Humphries et al., 1997; Kolset et al., 2002; Sampaio et al., 1992), indicating that these differences may reflect dynamism in the localization of HS-modifying enzymes along different Golgi cisternae according to the cellular context.

The redistribution of HS3ST5 along the Golgi was also observed in the presence of heparin. Shortly after treatment, the enzyme moved from cis to trans-Golgi, correlating with increased levels of trisulfated disaccharide in HS. These findings show that vesicular trafficking regulates the transport of HS-modifying enzymes throughout different Golgi compartments and that, eventually, this leads to the synthesis of different HS structures. Consequently, as shown previously for mucin O-glycosylation (Gill et al., 2010), depending on cellular context, substitution pattern may be changed due to redistribution of Golgi-resident proteins along the ER-Golgi pathway. This hypothesis was further confirmed by the analysis of HS-modifying enzymes, for which no significant changes in expression were observed. Also, an increase in sulfate levels could be explained by increased levels of PAPS synthase, which was not the case. Finally, it was shown that COPI subunits and Sec24 gene expression, which relates to COPII vesicles, changed, suggesting once again, that changes in cargo sorting, vesicular assembly and trafficking alter the dynamics of HS-modifying enzymes across the ER and Golgi, and that these changes lead to altered HS structure.

Collectively, the results show that cargo sorting, vesicular assembly and trafficking mediated by COPI and COPII regulate HS biosynthesis by controlling the spatial and temporal distribution of HS-modifying enzymes on different Golgi cisternae. The data demonstrate that cargo sorting, vesicular assembly and trafficking are critical prerequisites of the complete delineation of HS biosynthesis and the success of further structure and function studies. These findings also provide the basis for the development of bioengineered animal-free GAG-based therapeutics.

## Materials and Methods

### Reagents and antibodies

G418 disulfated salt solution was purchased from Sigma Aldrich (Saint Louis, MO, USA). Brefeldin A solution (1000X) (BFA) was obtained from Invitrogen (San Diego, CA, USA). Heparin (Hep) from porcine mucosa was a kind gift of Extrasul (Jaguapitã, PR, Brazil). H_2_ SO_4_ carrier free was purchased from National Centre for Nuclear Research Radioisotope POLATOM (Otwock, Poland). Mouse antibodies to HS3ST1 (B01P) and HS3ST3A1 (B01P) were obtained from Abnova (Taipei, Taiwan), antibodies to C5-epimerase and Golgin97 from Abcam (Cambridge, MA, USA) and antibody to NDST1 (M01) from Abgent (San Diego, CA, USA). Rabbit antibodies to anti-α-COP, β-COP and GM130 were purchased from Abcam, antibodies to COPII (Sec23) and HS3ST5 from Thermo Scientific (Rockford, IL, USA) and antibody to HS2ST (N-term) from Abgent. Goat antibody to GFP (I-16) was obtained from Santa Cruz Biotechnology (Dallas, TX, USA). Secondary antibodies conjugated to Alexa Fluor^®^ 488, Alexa Fluor^®^ 633 and Alexa Fluor^®^ 647 were purchased from Thermo Fisher Scientific. The information regarding all these antibodies are specified on Table S2.

### Cell culture

Endothelial cells derived from human umbilical vein endothelial cells (EC) (Takahashi et al., 1990) were maintained in F12 medium supplemented with 10% (v/v) fetal bovine serum (FBS, Cultilab, Campinas, Brazil), penicillin (100 U/mL) and streptomycin (100 µg/mL) (Gibco, CA, USA) at 37 °C in a humidified atmosphere of 2.5% CO_2_. At 80–85% confluence, the cells were detached with a solution of pancreatin (2.5%) diluted 1:10 (*v*/*v*) in EBSS, collected by centrifugation, suspended in F12 medium as described above (Buonassisi and Venter, 1976).

### Transfection and expression of HS3ST5 in culture

For cell transfection, EC cells were plated at 5 x 10^4^ cells per well (500 µL) in 24-well plates and transfected with 550 ng of cDNA coding HS3ST5, cloned into the vector pAcGFP-N1 (Clontech plasmid PT3716-5), using transfection FuGENE HD^®^ reagent at a ratio 5:1, according to the manufacturer’s instructions (Promega Corporation, WI, USA). The transfected cells (EC-HS3ST5) were cultured in the presence of G418 disulfated salt (0.5 µg/mL) and selected in accordance to the level of HS3ST5 expression.

### Immunofluorescence and confocal microscopy

EC and EC-HS3ST5 cells were seeded on 13 mm coverslips placed in 24-well plate (1 x 10^4^ cells/coverslip). After 4 days, the medium of transfected cells was removed and the cells treated with BFA (3 µg/mL) for 2 h or with heparin (20 µg/mL) from 1 to 4 h. The cells were then thrice washed with phosphate-buffered saline (PBS), fixed with 2% paraformaldehyde in PBS solution for 30 min at room temperature and washed with 0.1 M glycine. Afterwards, coverslips were sequentially incubated with blocking solution (0.02% saponin, 1% BSA in PBS solution, 30 min) and primary antibodies for 2 h. After washing with PBS, the cells were incubated with the appropriate fluorescent-labeled secondary antibodies for 1 h at room temperature. All antibodies were diluted in blocking solution. Once the first label was completed, labeling for the second protein was performed similarly. Nuclei were stained with 4′,6-diamidino-2-phenylindole (DAPI, Thermo Fischer Scientific, 1 μg/ml in blocking buffer). Lastly, coverslips were mounted on glass microscope slides using a mounting medium (Fluoromount-G, Birmingham, AL, USA) and fluorescence images were captured on a Leica TCS SP8 CARS confocal microscope (Wetzlar, Germany) with HC PL APO 63x/1.40 oil immersion objectives. The images represent the sum slides projections corresponding to the z-series of confocal stacks. The fluorescence images were quantified using the Leica LAS X Life Science software (Leica Microsystems) and colocalization intensity was expressed according to Pearson correlation values.

### Super-Resolution Ground State Depletion (SR-GSD) Microscopy

Transfected cells were seeded on 18 mm high precision round coverslips (Paul Marienfeld GmbH & Co. KG, Lauda-Königshofen, Germany) placed in 12-well plate (2 x 10^4^ cells/coverslip). After 3 days, the cells were thrice washed with iced PBS and fixed in two steps. Initially, cells were treated with buffer A (5 mM EGTA, 5 mM MgCl_2_, 5 mM glucose, 10 mM MES, 150 mM NaCl) containing 0.3% glutaraldehyde and 0.01% saponin for 2 min at room temperature and, then, with 0.5% glutaraldehyde diluted in buffer A for 10 min at room temperature. After washing, the cells were treated with 0.1% NaBH_4_ in PBS for 7 min at room temperature, washed and incubated with blocking solution (0.1% saponin, 5% FSB in PBS solution) for 1 h. The cells were then incubated with primary antibodies (1:50, diluted in PBS containing 0.1% saponin and 1% BSA) for 18 h at 4 °C. After washing, the cells were then incubated with appropriate fluorescent-labeled secondary antibodies (1:50, diluted in PBS containing 0.1% saponin and 1% BSA) for 90 min. Once the first label was completed, staining for the second protein was performed similarly. Lastly, the coverslips were mounted on depression slides containing embedding medium (70 mM β-mercapto-ethylamine in PBS solution). The images were captured on a Leica SR GSD 3D microscope (Wetzlar, Germany) equipped with a 160x high power super-resolution objective.

### Flow cytometry

EC and EC-HS3ST5 post-confluent cells (1 x 10^6^) were detached from the plate using EDTA 500 µM in PBS solution. The cells were washed with PBS, fixed with 2% paraformaldehyde in PBS for 30 min and, then, permeabilized with 0.01% saponin in PBS for 30 min. After, the cells were incubated with primary antibody for 2 h, followed by incubation with fluorescent-labeled secondary antibody for 40 min. The antibodies were diluted in PBS containing 1% BSA. Data were collected in the FACSCalibur flow cytometer (Becton Dickinson, Franklin Lakes, USA) and data analysis were performed using FlowJo v.10 software (Tree Star Inc, Ashand, USA).

### Composition analysis of HS disaccharides

Disaccharide composition analysis of HS extracted from transfected cells subjected to Hep stimulation was accomplished by enzymatic degradation followed by liquid chromatography (Vicente et al., 2015). Briefly, the transfected cells were subjected to metabolic labeling with carrier free [^35^S]-sulfate (150 μCi/ml) in serum-free F12 medium for 18 h at 37 °C in an atmosphere containing 2.5% CO_2_. Heparin (20 µg/mL) was added to medium from 1 to 4 h before completing the 18 hof radioactive sulfate incorporation. After labeling, the culture-conditioned medium was collected and the cells removed from the plate with 3.5 M urea in 25 mM Tris-HCl pH 8.0. Cell extract and medium were submitted to proteolysis with maxatase (proteolytic enzyme purified from *Bacillus subtilis*) (Biocon, Rio de Janeiro, RJ, Brazil) (4 mg/ml in 50 mM Tris-HCl, pH 8.0 containing 1.5 mM NaCl) at 60 °C. After proteolysis, nucleic acids and peptides were precipitated by the addition of 90% trichloroacetic acid (10% of sample volume). The GAGs present in the supernatant were precipitated with 3 volumes of iced methanol at −20 °C for 24 h. The precipitates formed (GAGs) were collected by centrifugation (4,000 rpm for 20 min at 4 °C), dried and suspended in 100 µL distillated water. The sulfated GAGs were identified and quantified by agarose gel electrophoresis in PDA buffer (0.05 M 1,3-diaminepropane acetate) (Dietrich and Dietrich, 1976). Lastly, 10,000 cpm of HS were incubated with 40 µL of each heparitinases I and II from *Flavobacterium heparinum* in 20 mM Tris-HCl, pH 7.4 containing 4 mM CaCl_2_ e 50 mM NaCl at 30 °C for 18h.The ^35^S-labeled degradation products were chromatographed in PhenoSphere™ 5 µM SAX (150 × 4.6 mm), using a NaCl gradient (0-0.3 M) for 30 min at a flow rate of 1 mL/min. Individual fractions (0.3 mL) were collected and counted on a Micro-Beta counter. The Δ-degradation products of HS were generated for three independent experiments and the products of digestion combined prior to analysis to allow detection. Therefore, the results represent an overall trend and were expressed as monosulfated, disulfated and trisulfated disaccharide groups.

### RNA extraction and real-time PCR

Total RNA was extracted from cultured cells using Trizol reagent (Invitrogen, Carlsbad, USA) according to the manufacturer’s instructions. RNA extraction was performed for wild type and transfected cells as well as to transfected cells subjected to heparin treatment (20 µg/mL). Reverse transcriptase reaction was performed from 2 µg of total RNA by using ImProm-II^TM^ Reverse Transcription System (Promega). Aliquots of cDNA obtained were amplified in PCR and quantitative real-time PCR reactions, using the primers described in Table S3. PCR reactions were performed using Master Mix (2X) (Promega) and carried out at an initial denaturation step of 95 °C for 2 min, followed by 35 cycles of denaturation at 95 °C for 30 s, annealing at 55 °C for 30 s and extension at 72 °C for 2 min, and final extension step at 72 °C for 5 min. The PCR products were analyzed on 1% agarose gels in TAE buffer at 100 V for 30 min. In addition, real-time PCR amplifications were performed using Maxima® SYBER Green Master Mix 2X (Fermentas, Waltham, MA, USA). The reactions were first subjected to an initial denaturation step at 95 °C for 10 min, followed by 40 cycles at 95 °C of 15 s (denaturation step) and at 60 °C for 1 min (annealing /extension steps). Melting curves were generated after the last amplification cycle to assess the specificity of the amplified products. The reactions were performed in triplicate on the 7500 Real Time PCR System (Applied Biosystems, Beverly, MA, USA). The relative expression levels of genes were calculated using the 2^-ΔCt^ method (Livak and Schmittgen, 2001). The transcript of ribosomal protein L13a (RPL13a) was used as a control to normalize the expression of target genes.

### Statistical analysis

Results were expressed as mean ± standard deviation of three independent experiments. Statistical analysis was determined by one-way analysis of variance (ANOVA) followed by Turkey test or Student’s *t*-tests. The statistical significance of differences was set at p < 0.05.

### Online supplemental material

Fig. S1 shows distribution prolife of HS-modifying enzymes (NDST1, C5-Epimerase, H3ST1 and HS3ST3A) in COPI and COPII vesicles. Fig. S2 shows fluorescently tagged HS3ST5 expression in transfected endothelial cells. Table S1 shows the relative amounts of monosulfated, disulfated and trisulfated disaccharides of HS extracted from transfected cells treated with heparin. Table S2 describes the set of antibodies used in immunofluorescence and flow cytometry assays. Table S3 shows the set of primers used in PCR and real-time PCR assays.

## Abbreviations

HS: heparan sulfate
Hep: heparin
GAG: glycosaminoglycan
EXT1: exostosin-1
EXT2: exostosin-2
HS3ST: heparan sulfate 3-O-sufotransferase
HS3ST1: heparan sulfate 3-O-sufotransferase 1
HS3ST2: heparan sulfate 3-O-sufotransferase 2
HS3ST3A: heparan sulfate 3-O-sufotransferase 3A
HS3ST5: heparan sulfate 3-O-sufotransferase 5
NDST1: N-deacetylase/N-sulfotransferase 1
HS2ST: heparan sulfate 2-O-sufotransferase
HS6ST: heparan sulfate 6-O-sufotransferase
BFA: brefeldin A
ER: endoplasmic reticulum
COPI: coatomer protein complex I
COPII: coatomer protein complex I
ERCIG: ER-Golgi intermediate compartment

## Acknowledgements

The work was supported by grants from Fapesp (Fundação de Amparo à Pesquisa do Estado de São Paulo, 2015/03964-6; 2017/14179-3); CAPES (Coordenação de Aperfeiçoamento de Pessoal de Nível Superior; 0599/2018) and CNPq (Conselho Nacional do Desenvolvimento Científico e Tecnológico; 442357/2014-1).

The authors declare no competing financial interests.

## Author Contributions

E.A.Y and M.A.L. contributed to the conception of the work; M.C.Z.M., H.B.N. and M.A.L. designed the experiment and analyzed the data; M.C.Z.M., P.D., C.M.V.P., L.P.B., R.P.C. and G.M.V. performed the experiments and analyzed the data; M.C.Z.M. drafted the work; H.B.N., E.A.Y and M.A.L. reviewed and edited the manuscript. All authors read and approved the manuscript.

