## Supplemental Material for "Heparan sulfate structure is influenced by the ER-Golgi dynamics of its modifying enzymes"

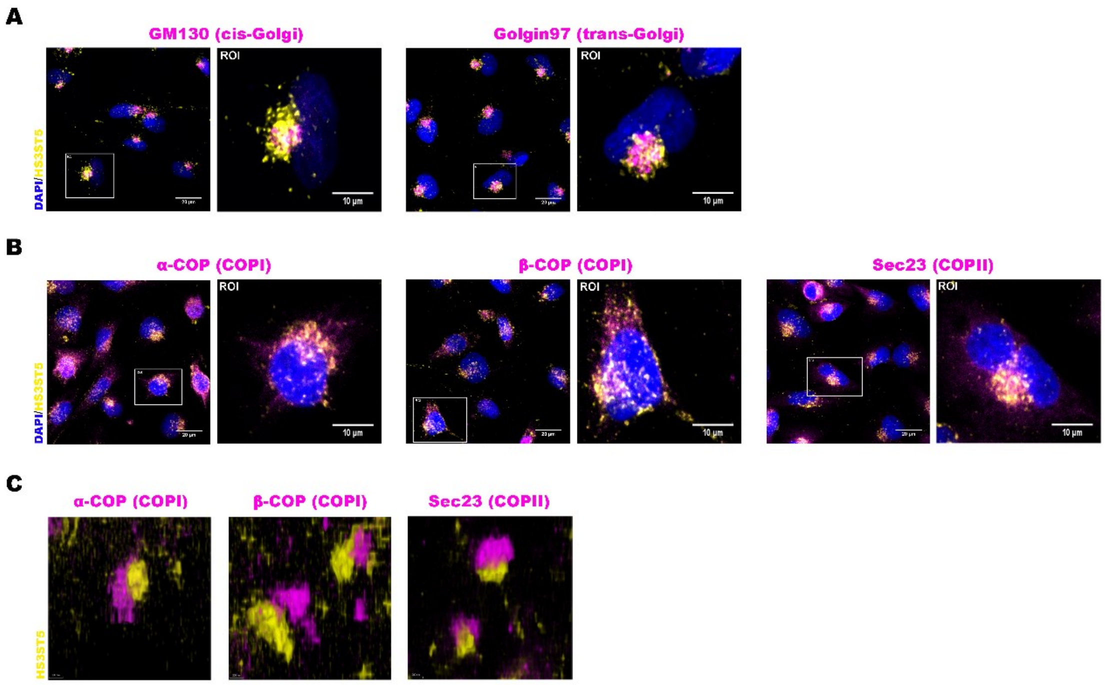


**Figure 1. Subcellular localization of fluorescent HS3ST5.**

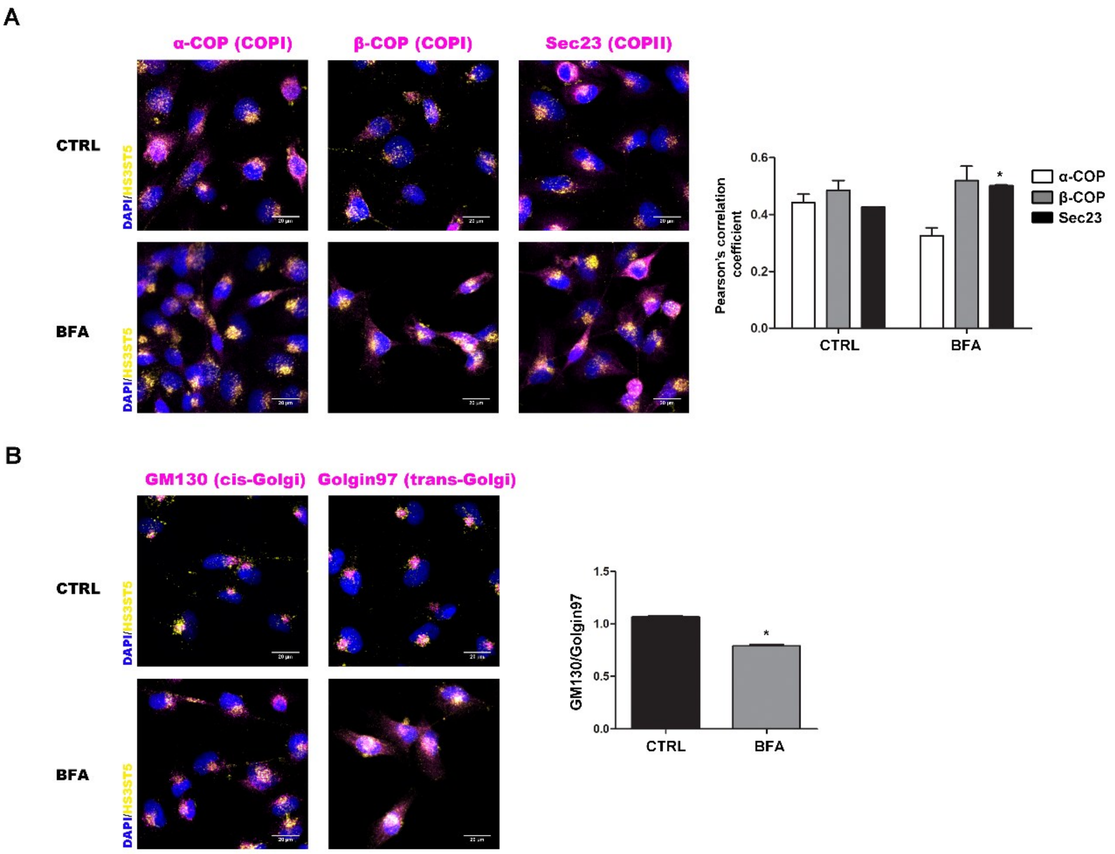


**Figure 2. Distribution profile of HS3ST5 in secretory pathway in the presence of brefeldin A.**

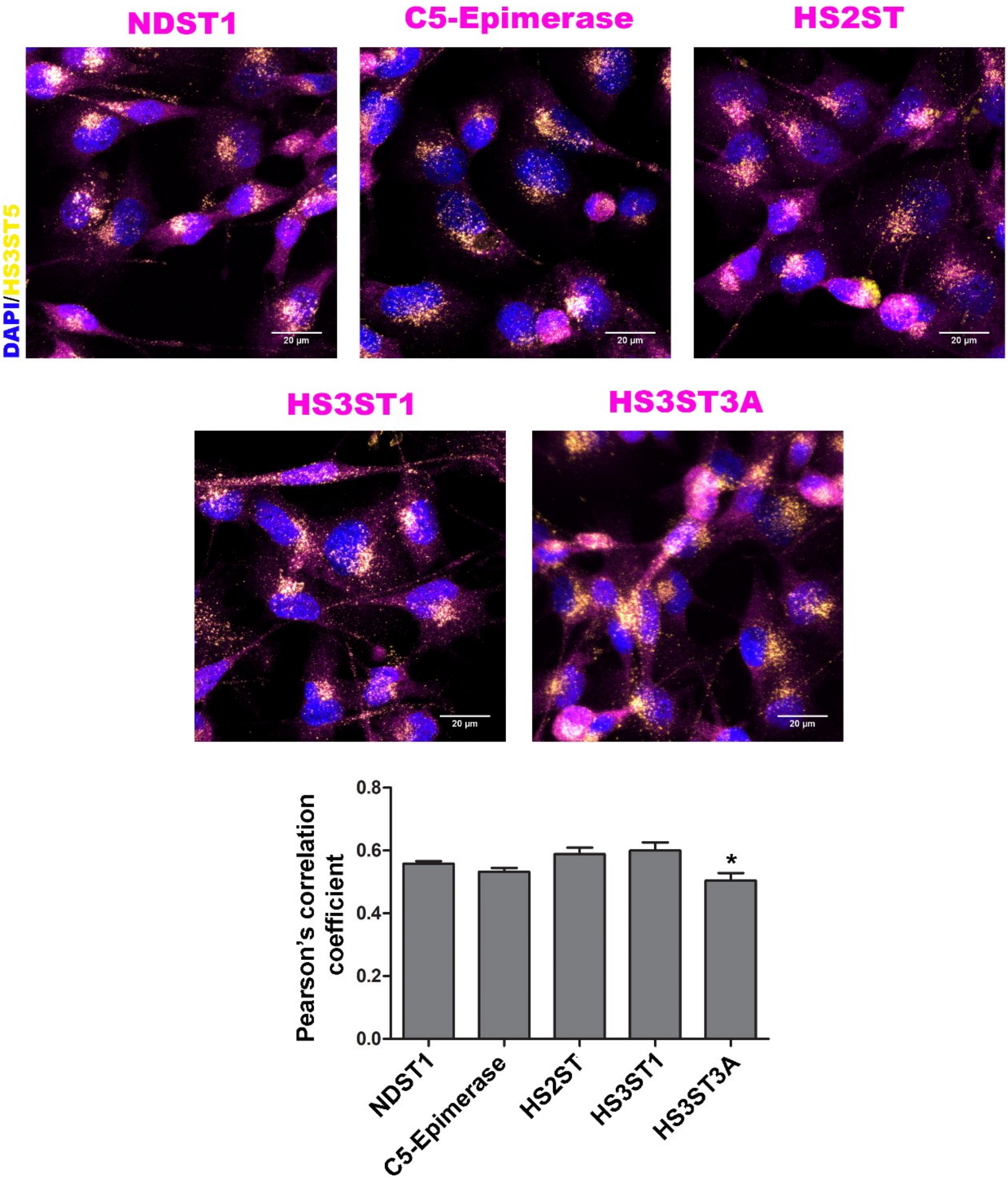


**Figure 3. Distribution profile of HS-modifying enzymes following BFA treatment.**

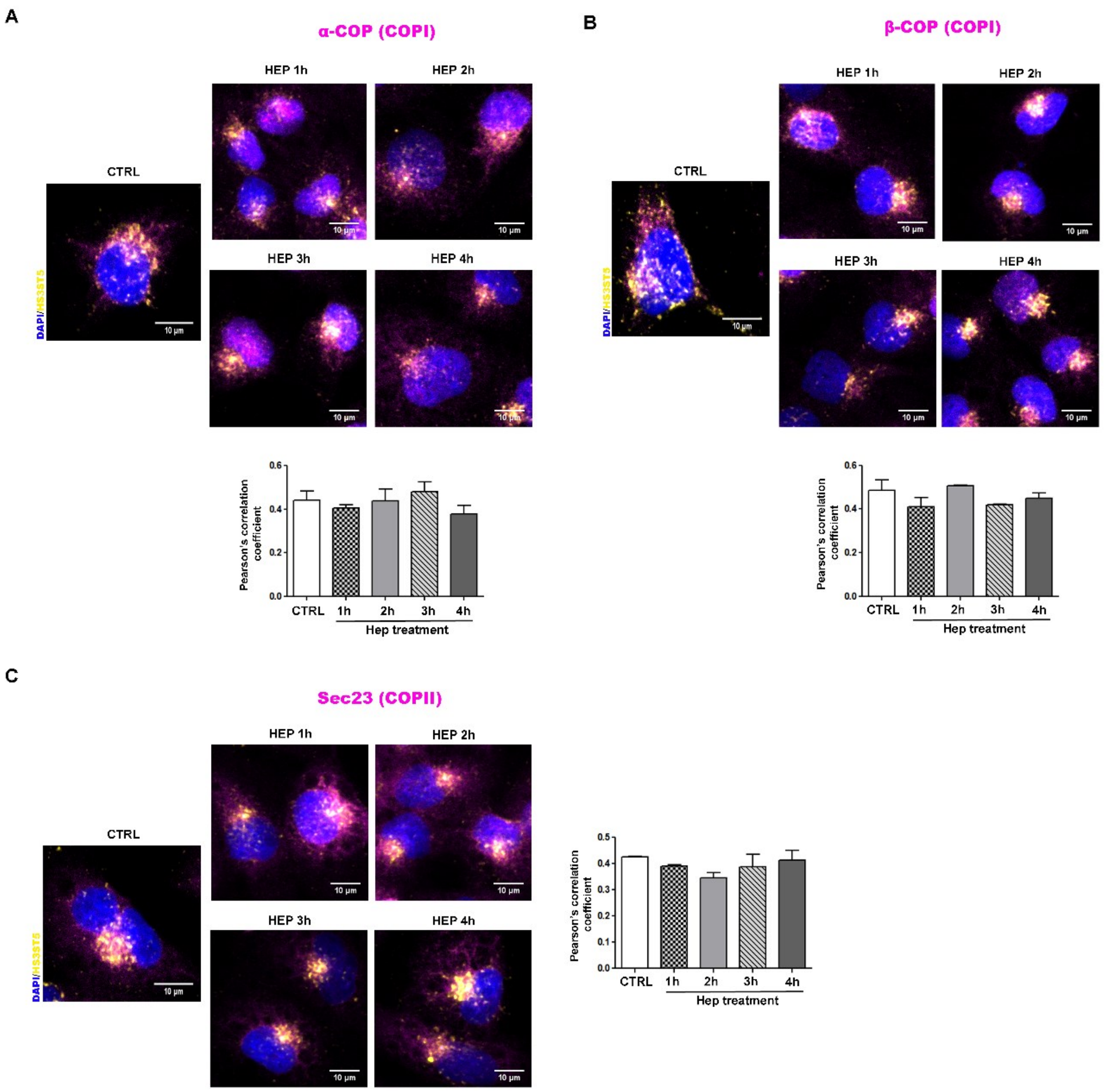


**Figure 4. Distribution profile of HS3ST5 in coated vesicles in the presence of heparin.**

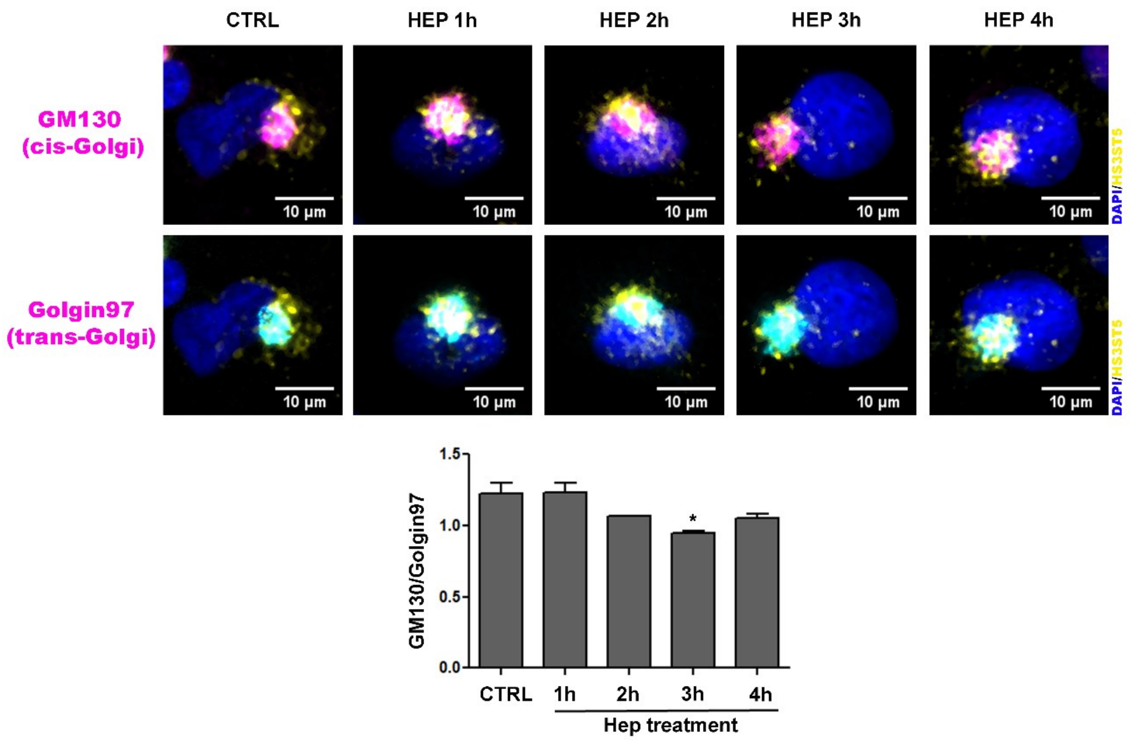


**Figure 5. Distribution profile of HS3ST5 in Golgi apparatus following heparin treatment.**

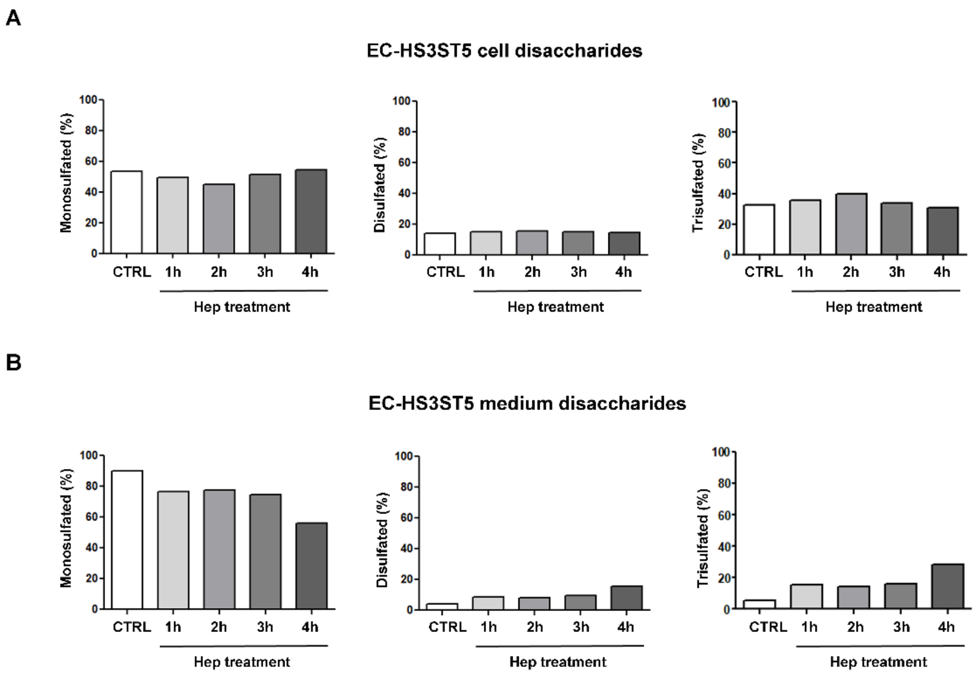


**Figure 6. Disaccharide composition of HS extracted from EC-HS3ST5 cells treated with heparin.**

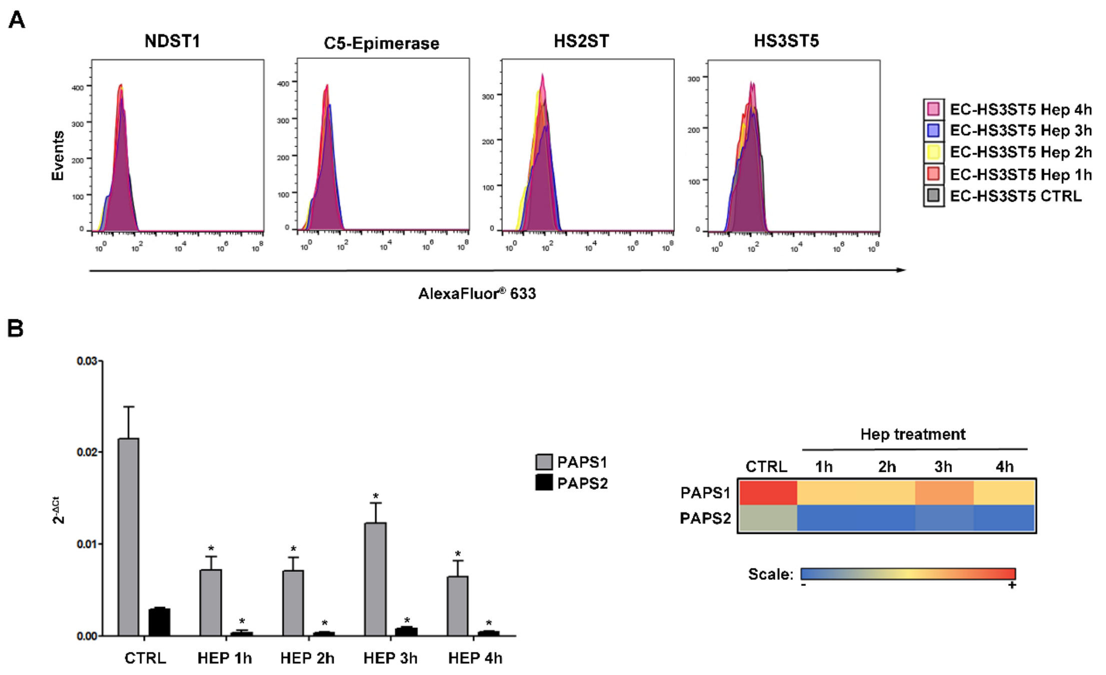


**Figure 7. Protein and gene expression of components of HS biosynthesis in presence of heparin.**

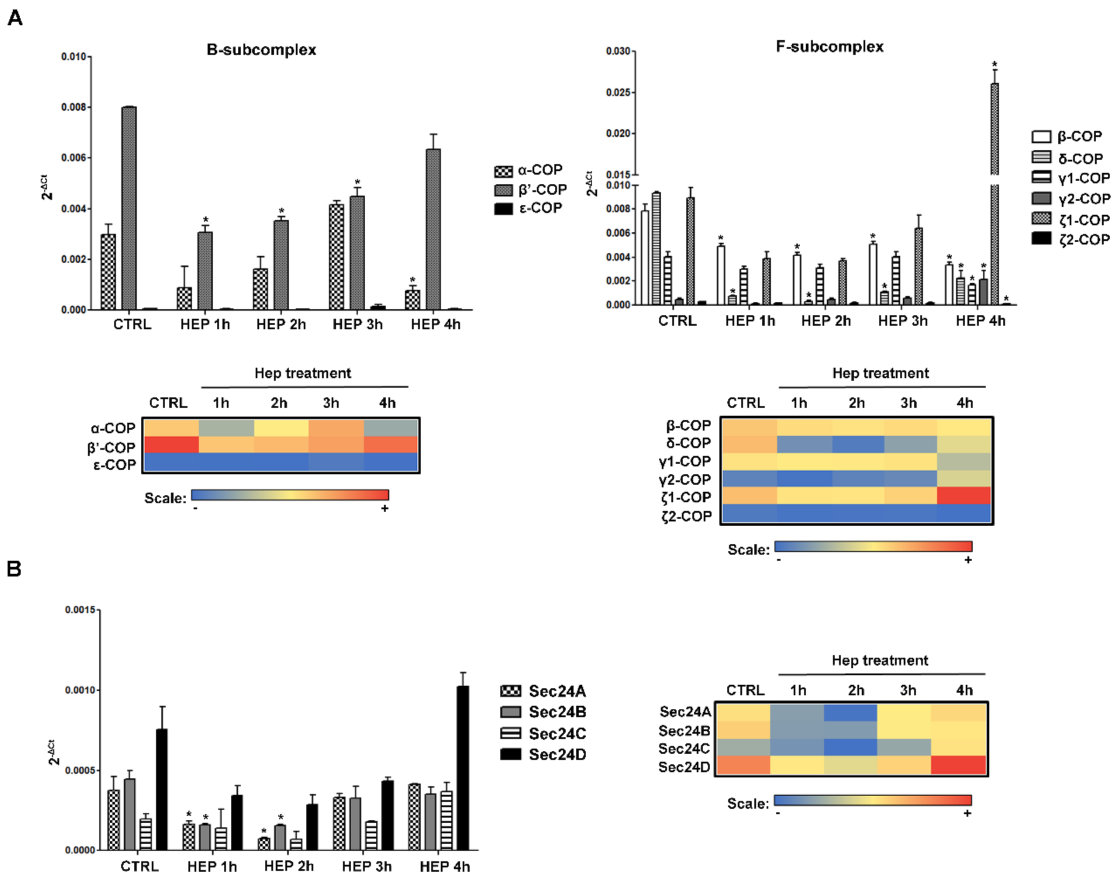


**Figure 8. Gene expression of coated vesicles subunits in the presence of heparin.**
